# Low nutrient levels reduce the fitness cost of *MexCD-OprJ* efflux pump overexpression in ciprofloxacin-resistant *Pseudomonas aeruginosa*

**DOI:** 10.1101/298471

**Authors:** Wenfang Lin, Kun Wan, Jie Zeng, Jingjing Li, Xi Li, Xin Yu

## Abstract

The long-term persistence of antibiotic resistance in the environment is a public health concern. Expression of an efflux pump, an important mechanism of resistance to antibiotics, is usually associated with a fitness cost in bacteria. In this study, we aimed to determine why antibiotic resistance conferred by overexpression of an efflux pump persists in environments such as drinking and source water in which antibiotic selective pressure may be very low or even absent. Competition experiments between wild-type *Pseudomonas aeruginosa* and ciprofloxacin-resistant mutants revealed that the fitness cost of ciprofloxacin resistance (strains cip_1, cip_2, and cip_3) significantly decreased (P < 0.05) under low-nutrient (0.5 mg/l total organic carbon (TOC)) relative to high-nutrient (500 mg/l TOC) conditions. Mechanisms underlying this fitness cost were analyzed. *MexD* gene expression in resistant bacteria (cip_3 strain) was significantly lower (P < 0.05) in low-nutrient conditions, with 10 mg/l TOC (8.01 ± 0.82-fold), than in high-nutrient conditions, with 500 mg/l TOC (48.89 ± 4.16-fold). Moreover, *rpoS* gene expression in resistant bacteria (1.36 ± 0.13-fold) was significantly lower (P < 0.05) than that in the wild-type strain (2.78 ± 0.29-fold) under low-nutrient conditions (10 mg/l TOC), suggesting a growth advantage. Furthermore, the difference in metabolic activity between the two competing strains was significantly smaller (P < 0.05) in low-nutrient conditions (5 and 0.5 mg/l TOC). These results suggest that nutrient levels are a key factor in determining the persistence and spread of antibiotic resistance conferred by efflux pumps in the natural environment with trace amounts or no antibiotics.

**Importance:** The widespread of antibiotic resistance has led to an increasing concern about the environmental and public health risks. Mechanisms associated with antibiotic resistance including efflux pumps often increase bacterial fitness cost. Our study showed that the fitness cost of ciprofloxacin resistance conferred by overexpression of *MexCD-OprJ* efflux pump significantly decreased under low-nutrient relative to high-nutrient conditions. The significance of our research is to reveal that nutrient levels are key factor in determining the persistence of antibiotic resistance conferred by efflux pumps under conditions with trace amounts or no antibiotics, which can be mediated by some mechanisms including *MexD* gene expression, SOURs differences, and *rpoS* gene regulation.

## 1 Introduction

Continuing imprudent use of antibiotics has enriched antibiotic resistant bacteria in the environment. The spread of antibiotic-resistant pathogens has become a serious problem for human health around the world. Generally, bacterial antibiotic resistance is achieved through four main mechanisms (1): a reduction in outer membrane impermeability (2), enzymatic inactivation (3), target alterations (4), and active efflux of antibiotics (5). Among these, rapid efflux of antibiotics from cells is thought to play a key role in the intrinsic resistance of clinically important bacteria, including *Pseudomonas aeruginosa*, *Escherichia coli*, and *Salmonella* Typhimurium (5, 6). To date, five families of bacterial efflux systems have been identified (5, 7): the resistance-nodulation-division family (RND), major facilitator (MF) family, multidrug and toxic efflux (MATE) family, small multidrug resistance (SMR) family, and ATP-binding cassette (ABC) family.

*P. aeruginosa* is a ubiquitous opportunistic human pathogen with high levels of antibiotic resistance. Four multidrug efflux systems belonging to the RND family have been well characterized in *P*. *aeruginosa*, including *MexAB-OprM*, *MexCD-OprJ*, *MexEF-OprN*, and *MexXY-OprM* (8-10). Efflux pump genes are often part of an operon, with a regulatory gene that controls expression. For example, *MexCD-OprJ* has been shown to confer resistance to fluoroquinolones such as ciprofloxacin. *NfxB* is a transcriptional regulator that tightly represses the expression of *MexCD-OprJ* in wild-type strains. Overexpression of efflux pumps can result from mutations in repressor genes. In resistant *P*. *aeruginosa*, a mutation in *nfxB* leads to derepression of *MexCD-OprJ*, causing high levels of ciprofloxacin resistance (11).

It is generally accepted that the acquisition of antibiotic resistance imposes fitness cost on bacteria (12). Resistance acquired through mutation of elements with important physiological roles imposes a metabolism burden on bacteria. Consequently, wild-type bacteria might outcompete resistant bacteria in the absence of antibiotic selective pressure. For example, *nfxB* mutants have only rarely been detected in a clinical setting since they were first described by Hirai two decades ago (13). An attractive hypothesis is that these bacteria were avirulent because of impaired fitness. Additionally, some previous studies have suggested that *nfxB* mutations in *P. aeruginosa* impair bacterial growth, all forms of motility (swimming, swarming, and twitching), and metabolic products such as siderophores, rhamnolipid, secreted protease, and pyocyanin, or that these mutations have led to specific changes in bacterial physiology (11, 14, 15).

The fitness cost of antibiotic resistance is strongly dependent on experimental conditions. For example, some resistance mutations have shown no cost in laboratory medium but a high cost in mice (16). Additionally, environmental factors such as temperature and resource availability affect the fitness cost of rifampicin resistance mutations (17). However, few studies have examined the contributions of efflux pumps to the fitness cost of bacterial antibiotic resistance. Overexpression of efflux pumps is known to consume a lot of energy, which may present a general burden to bacteria in the absence of antibiotics. In addition, bacteria utilize carbon compounds as an important energy source. Thus, we hypothesized that the availability of a carbon source would determine the fitness cost of antibiotic resistance conferred by overexpression of efflux pump in low-nutrient environments such as drinking or source water.

In this study, the effect of the environmental availability of nutrients on the fitness cost of bacterial antibiotic resistance conferred by overexpression of efflux pumps was investigated. The important opportunistic pathogen *P*. *aeruginosa* and its *MexCD-OprJ* efflux pump were the focus of this study. We aimed to identify whether *MexCD-OprJ* overexpression imposed a fitness cost on *P*. *aeruginosa* in different nutrient levels and to analyze the mechanisms underlying any cost.

## 2. Materials and methods

### 2.1 Strains and growth conditions

Bacterial strains used in this study are described in Table 1. Ciprofloxacin resistance in *P. aeruginosa* PAO1 were induced by the mutagenic disinfection byproduct dichloroacetonitrile (DCAN) (18). One ciprofloxacin-resistant strain (cip_1) had mutations in both the *nfx*B gene (L83P) and *parC* gene (S65F). In addition, a mutation was identified in the *nfx*B gene of the cip_3 (L14Q) and cip_4 (L83P) strains. Moreover, an insertion mutation at nt 32 occurred in the cip_2 *nfx*B gene. These mutants showed increased resistance to ciprofloxacin. The minimal inhibitory concentration (MIC) of ciprofloxacin in these mutants was 2 μg/ml, whereas the MIC in wild-type *P. aeruginosa* PAO1 was 0.125 μg/ml. Bacteria cells were grown routinely in Luria-Bertani (LB) broth, with shaking at 200 rpm, 37 °C, and 0.5 × MIC or without ciprofloxacin.

**Table 1:** Bacterial strains used in this study.

| Strains | Mutation in *nfxB* | Mutation in *parC* | MICs (μg/ml) |
| --- | --- | --- | --- |
| cip_1 | L83P(CTG→CCG) | S65F(TCC→TTC) | 2 |
| cip_2 | 32-nt insert A | — | 2 |
| cip_3 | L14Q(CTG→CAG) | — | 2 |
| cip_4 | L83P(CTG→CCG) | — | 2 |
| *P. aeruginosa* PAO1 | — | — | 0.125 |

### 2.2 Fitness cost measurement

Fitness costs were determined by competition experiments between ciprofloxacin-resistant mutant and wild-type strains (19, 20). Briefly, overnight cultures of resistant and wild-type cells were washed twice with sterile saline solution. Resistant cultures were mixed with wild-type cultures at a ratio of 1:1, and 1.5 μl of this mixture was used to inoculate 15 ml of fresh artificial synthetic wastewater (SW) medium (21) under different nutrient conditions (total organic carbon [TOC] concentrations of 500, 50, 5, 0.5, and 0.05 mg/l) in the absence of antibiotics and at a starting cell density of 10^5^ CFU/ml. Additionally, the SW medium was sterilized by pasteurization (70 °C for 30 min) to avoid potentially damaging components in the medium. Glucose was used as the sole carbon source in the SW medium for bacterial growth. Preliminary data revealed that *P. aeruginosa* PAO1 reached the stationary phase after 12 h in SW medium. Therefore, every 12 h, 1.5 ml of the cultures were inoculated into 15 ml of fresh SW medium for growth (15).

To determine the number of viable cells, the cultures were serially diluted by 1:10 in saline solution, and suitable dilutions were plated every 12 h on antibiotic-free LB agar to count the total number of colonies. In parallel, plates containing antibiotics (1 mg/l ciprofloxacin) allowed the growth of mutant strains. The number of wild-type cells was calculated as the total number of bacterial cells minus the number of antibiotic-resistant cells. Competition experiments were performed in triplicate. One mutant (cip_3) was selected for application in the following mechanistic analysis.

### 2.3 Mechanistic analysis

#### 2.3.1 Quantitation of *MexD* gene expression by Reverse transcription quantitative real-time PCR (RT-qPCR)

To investigate the effect of an efflux pump on the fitness of a resistance mutant under different nutrient conditions, *MexD* gene expression was quantified by RT-qPCR. The Cip_3 strain was grown in LB broth containing ciprofloxacin antibiotic overnight at 37 °C. Afterwards, the cultures were washed twice with saline solution and then re-suspended in 1 ml of the same buffer (optical density at 600 nm, 0.3; ~1 × 10^9^ CFU/ml). A 20-μl aliquot of the pre-culture was inoculated into 200 ml of fresh SW medium containing 500 mg/l and 10 mg/l TOC (initial cell density, ~10^5^ CFU/ml), and shaken at 37 °C. Simultaneously, *MexD* gene expression in wild-type *P. aeruginosa* PAO1 was quantitated as a control.

The cells were harvested in the stationary phase by centrifugation (7800 rpm for 20 min). Total RNA was extracted from cell pellets using an RNA isolation kit (TransGene, China) according to the manufacture. After that, RNA was converted into cDNA using a cDNA synthesis kit (TransGene, China) according to the manufacturer’s instructions to avoid RNA degradation. Primers used in this study are shown in Table S1. Quantitative RT-qPCR of the cDNA was performed on an ABI 7300 detection system in 20-μl reaction mixtures containing 100 ng isolated RNA, 200 nM each of the two primers, and 10 µl 2× SYBR green PCR mixture. After an initial 2-min incubation at 95 °C, the reaction was subjected to 35 cycles of 95 °C for 40 s, 60 °C for 30 s, and 72 °C for 40 s, and then a final 7-min incubation at 72 °C. A constitutively expressed gene (16S rRNA) was used as a control to normalize the results, and the amount of each RNA was calculated following the 2^△△-ct^ method (22).

#### 2.3.2 Quantitation of *rpoS* gene expression by RT-qPCR

To quantify *rpoS* gene expression, the bacterial strain (cip_3) was grown to the early stationary phase in SW medium supplemented with 500 mg/l and 10 mg/l TOC and then harvested by centrifugation. Total RNA extraction and RT-qPCR were performed as described in 2.3.1.

#### 2.3.3 Specific oxygen uptake rate (SOUR) measurement

SOUR was measured to evaluate differences in metabolic activity between resistant and wild-type cells under different nutrient conditions. One mutant (cip_3) was selected as an example. The bacteria were cultured in LB broth at 37 °C overnight and washed twice with saline solution by centrifugation at 7800 rpm for 10 min. The SW media containing 500 mg/l, 5 mg/l, and 0.5 mg/l TOC was aerated until it was saturated with dissolved oxygen (DO), and then the cell pellets were resuspended. The cell density was approximately 10^10^ CFU/ml, and the oxygen concentration decreased with respiration of the bacteria. DO concentrations were recorded using a DO meter (Germany, Multi 3420). The slope in the DO concentration versus time was used to obtain SOUR values. All SOUR assays were conducted in triplicate.

### 2.4 Data analysis

#### 2.4.1 Determination of bacterial fitness

The fitness cost of an efflux pump in a resistance mutant was evaluated by determining two values: the ratios of the competing strains (*u*) and their fitness (*fit_t_*). *u* is a denary logarithmic transformation of the ratios of the numbers of ciprofloxacin-resistant and wild-type cells at time *t*. To eliminate the effect of initial cell density, ratios of the cells numbers at time *t* were normalized to that at time *t_0_* as follows:

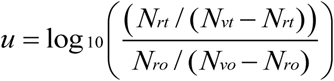

where (*N_rt_* and *N_vt_* − *N_rt_*) and (*N_ro_* and *N_vo_* − *N_ro_*) denote the absolute number of ciprofloxacin-resistant and wild-type cells at time *t* and *t_0_*, respectively. *u* is equal to 0 if there is no difference in fitness cost between the competing strains, *u* is positive if resistance reduces the bacterial fitness cost, and *u* is negative if resistance increases the bacterial fitness cost.

In addition, the bacterial fitness (*fit_t_*) of the two competing strains was calculated from the quotient of the growth rates of the competing strains at time *t* and the preceding time point *t-1*. This quotient was standardized with the exponent 1/*n*. *n* is the number of generations of bacterial growth from *t-1* to *t*. *fit_t_* is 1 plus the natural logarithmic transformation of the quotient using the following function (19):

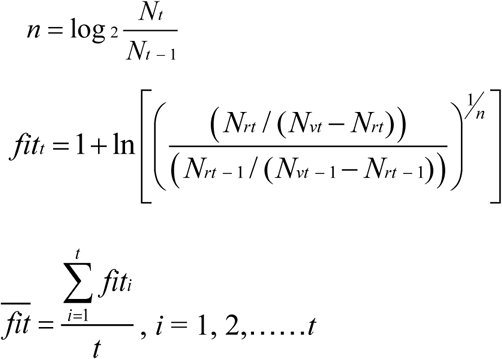

Furthermore, 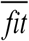 denotes the average fitness (*fit_t_*). *fit_t_* or 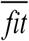is equal to 1 if there is no difference in fitness cost between the competing strains, *fit_t_* or 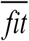 is above 1 if resistance reduces the bacterial fitness cost, and *fit_t_* or 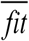 is below 1 if resistance increases the bacterial fitness cost.

#### 2.4.2 RT-qPCR

The relative expressions of *MexD* and *rpoS* gene were estimated by RT-qPCR. Additionally, the 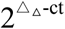 method was used to calculated the qPCR data as follows:

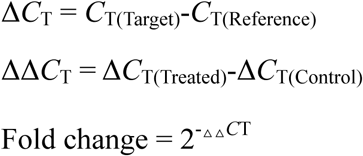

Where Δ*C*_T_ is the difference in *C*_T_ values between the target gene (*MexD* or *rpoS*) and the endogenous reference gene (16S rRNA) for each sample, and ΔΔ*C*_T_ is the difference in the Δ*C*_T_ values for the two samples (treated and control). The treated sample is the bacterial strain grown in low-nutrient medium (5 mg/l TOC), and the control samples is the bacterial strain grown in high-nutrient medium (500 mg/l). The fold change in relative expression levels was calculated using the 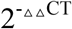 method. Genes expression levels were considered elevated in the treatment groups when the fold change was >1.

### 2.4.3 Statistical analysis

Analysis of variance (ANOVA) and Kruskal-Wallis tests were used to determine significant differences in bacterial fitness (*fit_t_*) and SOUR under different nutrient conditions. Additionally, differences in *MexD* and *rpoS* gene expression levels under high and low-nutrient conditions were analyzed using Student’s t-test. Results were significant at the 95% level (p < 0.05).

## 3. Results and discussion

### 3.1 Growth curves under different nutrient conditions

The wild-type strain followed different growth curves than the ciprofloxacin-resistant strains (Fig. S1). As shown in Fig. S1, the wild-type strain grew to approximately 10^7^–10^8^ CFU/ml 24 h after inoculation in the medium containing 500, 50, and 5 mg/l TOC. However, bacterial growth became unstable under low-nutrient conditions. It fluctuated around 10^6^ CFU/ml in the medium containing 0.5 mg/l TOC and at about 10^5^–10^6^ CFU/ml in the medium supplemented with 0.05 mg/l TOC. The cell concentrations depended on the available nutrients in the medium because the carbon source provided energy for microbial metabolism.

In contrast, the concentrations of the ciprofloxacin-resistant strains showed a downward trend under high-nutrient conditions (Fig. 1). The cell density reached 10^6^– 10^7^ CFU/ml 24 h after inoculation in the medium containing 50 mg/l TOC, but it decreased rapidly to 10^5^ CFU/ml (cip_1), 10^1^ CFU/ml (cip_3), and 10^5^ CFU/ml (cip_4). Similarly, the cell density decreased from 10^6^ CFU/ml to 10^2^ CFU/ml (cip_1), 10^1^ CFU/ml (cip_3), and 10^5^ CFU/ml (cip_4) under high-nutrient conditions with 500 mg/l TOC (Fig. 1).

**Fig. 1.**
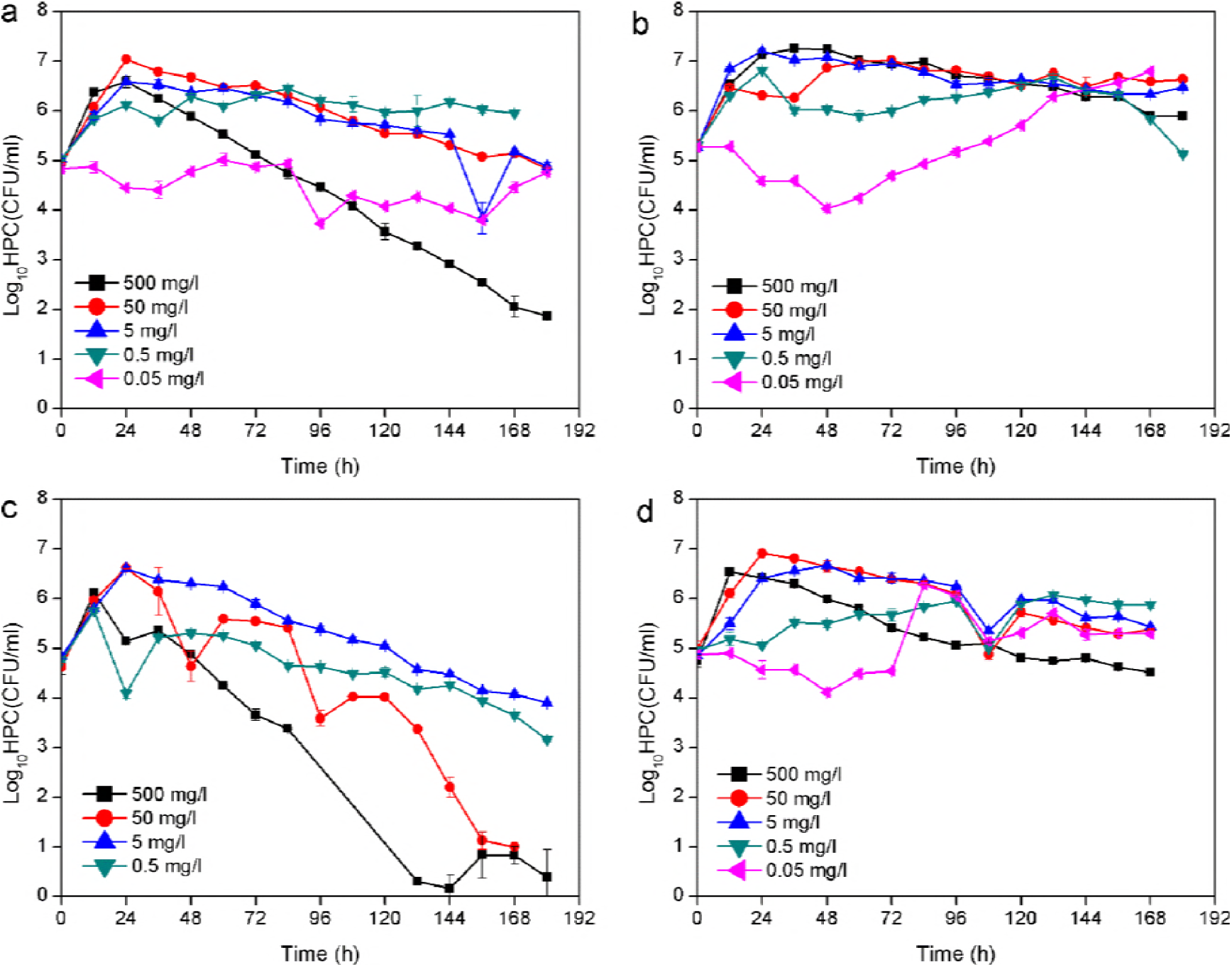
Growth curves of ciprofloxacin-resistant *P. aeruginosa* PAO1 during a competition experiment. (a) cip_1, (b) cip_2, (c) cip_3, (d) cip_4.

However, growth of the resistant mutants was more stable under low-nutrient conditions. For instance, cell concentrations were maintained at 10^5^–10^6^ CFU/ml in the presence of 0.5 mg/l TOC (cip_1 and cip_4). Similarly, the density of resistant strains was maintained at approximately 10^4^–10^5^ CFU/ml in the medium supplemented with 0.05 mg/l TOC (cip_1). Additionally, the cell density of cip_3 decreased in all nutrient conditions (500, 50, 5, and 0.5 mg/l TOC), but it showed a slower decrease at lower nutrient levels. Moreover, the cip_2 resistance mutant remained at 10^6^ CFU/ml in the medium supplemented with 0.5 mg/l TOC and fluctuated under low-nutrient conditions (0.05 mg/l TOC), although it was reduced only minimally, from 10^7^ CFU/ml to 10^6^ CFU/ml, in high-nutrient concentrations (500, 50, and 5 mg/l TOC). These results revealed that nutrient levels affected the growth of ciprofloxacin resistance mutant cells.

### 3.2 Ratio of cell numbers and fitness costs for the two competing strains

The ratios of two competing strains (*u*) decreased under high-nutrient conditions (Fig. 2). Using cip_2 as an example (Fig. 2b), the value of *u* decreased to −2 in the medium containing 500 and 50 mg/l TOC. Additionally, this value gradually decreased to −1 at a nutrition level of 5 mg/l TOC. This result indicated that the wild-type strain outcompeted the *nfxB* mutant that was resistant to ciprofloxacin. The acquisition of antibiotic resistance is generally assumed to represent an extra metabolic burden that affects bacterial fitness (23). However, the value of *u* fluctuated around 0 in the medium supplemented with 0.5 mg/l TOC. Furthermore, this value increased to 1 at 0.05 mg/l TOC. Similar trends were also observed for cip_1, cip_3, and cip4. These data indicate that the number of ciprofloxacin-resistant mutant cells was reduced less or even outcompeted by the wild-type strain under low-nutrient conditions.

**Fig. 2.**
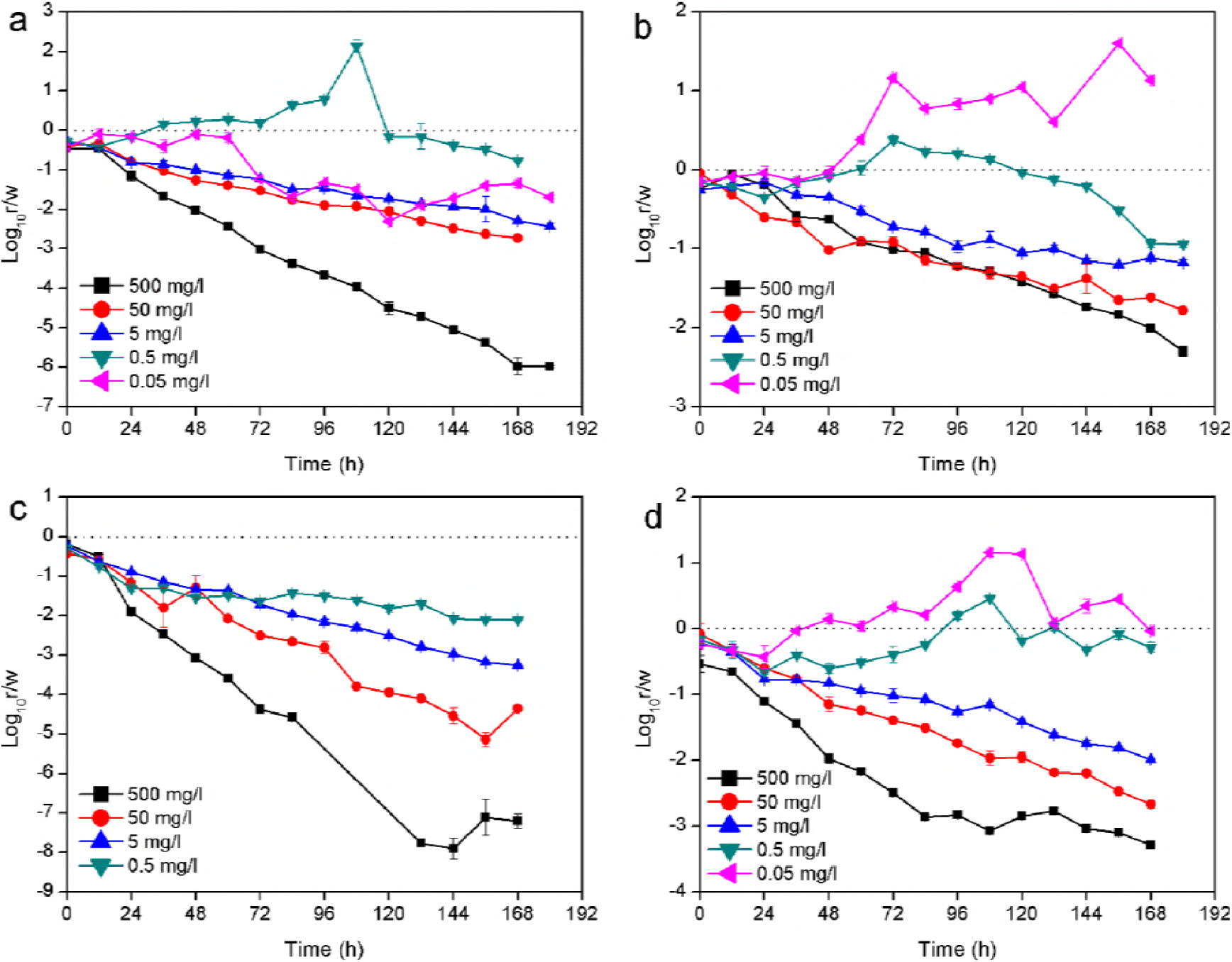
Change in the ratio of the number of ciprofloxacin-resistant *P. aeruginosa* PAO1 cells and cells of the wild-type strain. (a) cip_1, (b) cip_2, (c) cip_3, (d) cip_4.

**Fig. 3.**
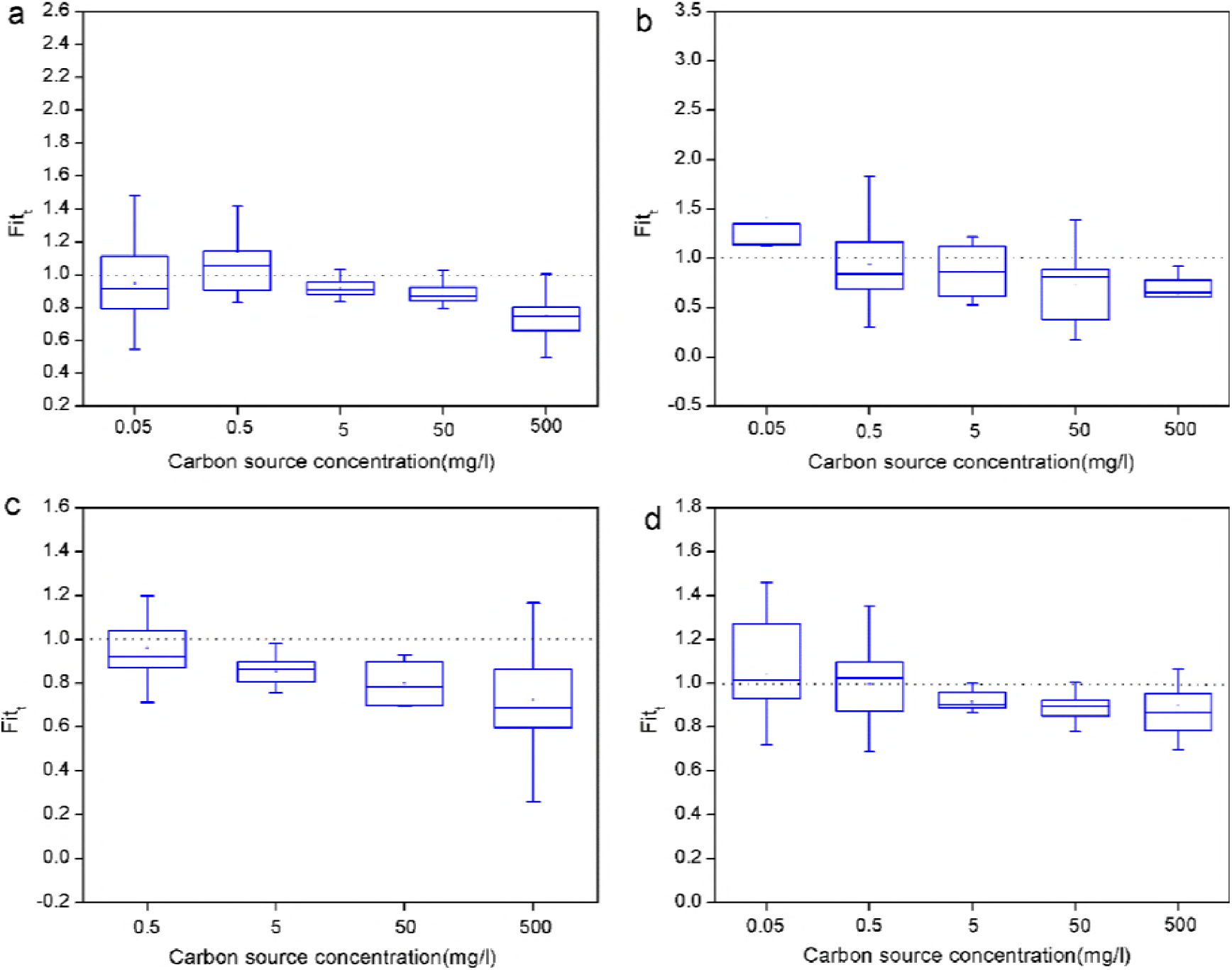
Box plots showing change in bacterial relative fitness (*fit_t_*) at different nutrient levels, (a) cip_1, (b) cip_2, (c) cip_3, (d) cip_4. Symbols indicate the following: box, 25^th^ to 75^th^ percentile; horizontal line, median; square, mean value; and whiskers, 10^th^ and 90^th^ percentile.

In addition, the relative fitness (*fit_t_*) of the bacteria was also calculated (Fig. S2 and 3). The values of *fit_t_* were found to be below 1 more frequently at higher nutrient levels (500, 50, and 5 mg/l TOC). However, the *fit_t_* values were close to or above 1 for strains grown in medium with low nutrient levels (0.5 and 0.05 mg/l TOC). Next, the average fitness 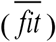 was calculated to more directly compare the fitness cost of resistant strains under different nutrient levels (Fig. 4). The average fitness 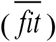 of the four resistant mutants (cip_1, cip_2, cip_3, and cip_4) were all below 1 under higher nutrient conditions (5, 50, and 500 mg/l TOC), suggesting considerable fitness costs in the resistant bacteria. This result was in line with those of previous studies, in which *nfxB* mutants conferring resistance to ciprofloxacin were shown to impair fitness, expressed as a reduced growth rate or altered virulence and metabolite production (11, 14).

**Fig. 4.**
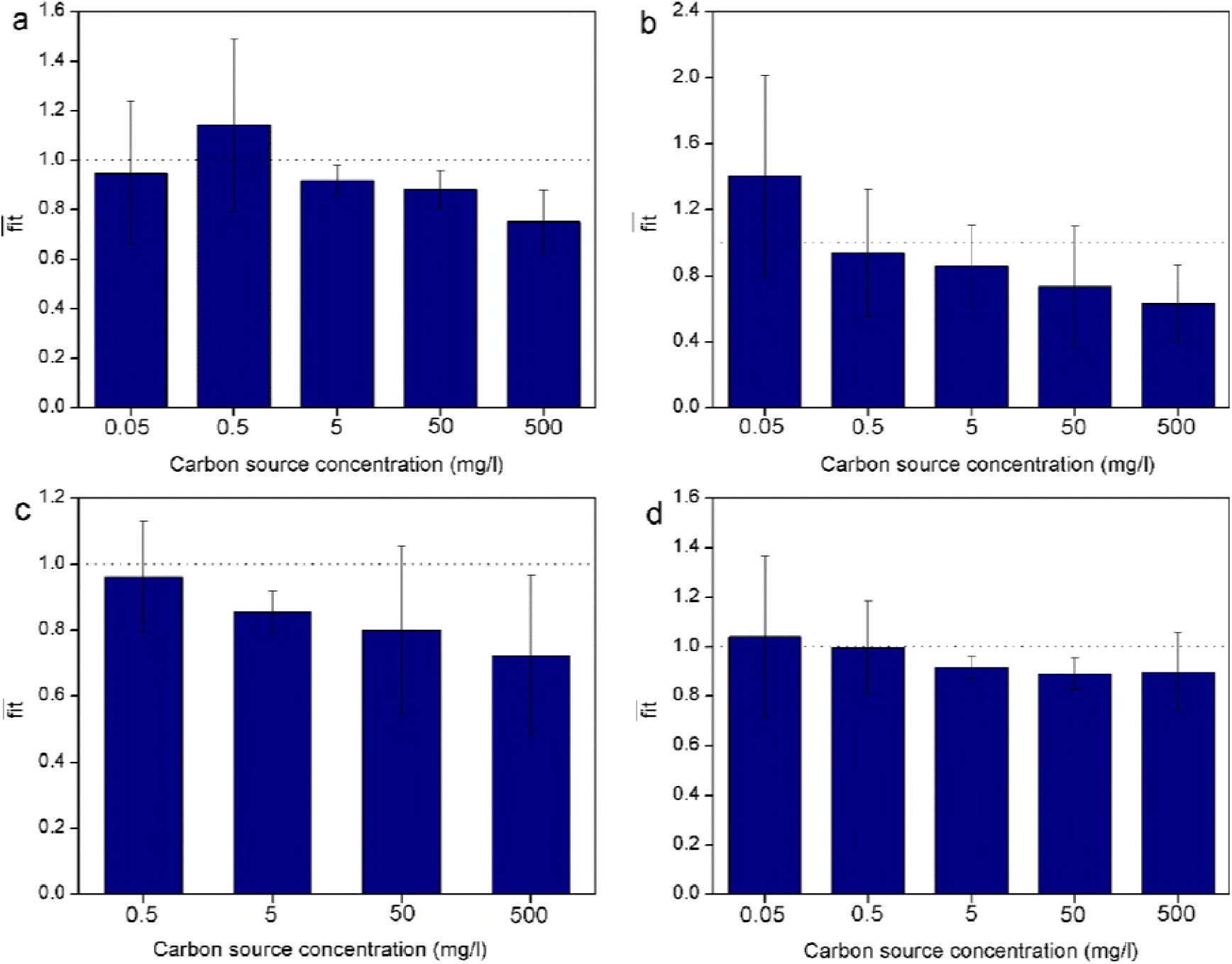
Change in the average bacterial fitness 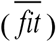 at different nutrient levels. (a) cip_1, (b) cip_2, (c) cip_3, (d) cip_4.

However, the fitness values increased with decreasing nutrient levels. Specifically, this value was above 1 in for cip_2 and cip_4 grown in medium containing 0.05 mg/l TOC (1.41 ± 0.61 and (1.04 ± 0.32, respectively). Additionally, the value of 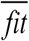 for cip_1 and cip_4 reached nearly 1 at 0.5 mg/l TOC (1.14 ± 0.35 and 0.99 ± 0.19, respectively). These results suggested that the fitness cost of the *nfxB* mutants was reduced in low-nutrient conditions. Furthermore, the values of 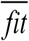 for cip_1, cip_2, and cip_3 under low-nutrient conditions (0.5 mg/l) were significantly larger (P < 0.05) than under high-nutrient conditions (500 mg/l TOC). Furthermore, this value for cip_2 was also significantly larger (P < 0.05) at low nutrient levels (0.05 mg/l TOC) than at high nutrient levels (500 mg/l TOC). For cip_4, no significant differences in the 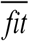 values were observed in different nutrient conditions, but an increasing trend with decreasing nutrient levels was apparent. Consequently, the increased numbers of ciprofloxacin-resistant cells under low-nutrient conditions (Fig. 1) can be explained by a decreasing fitness cost, and possible underlying mechanisms were explored in the following experiments.

## 3.3 Mechanisms analysis

### 3.3.1 Efflux pump *MexD* gene expression

As mentioned above, mutations in *nfx*B (L83P, L14Q, and L83P) caused ciprofloxacin resistance in the cip_1, cip_3, and cip_4 strains, respectively. Additionally, an insertion mutation at nt 32 was present in *nfx*B in the cip_2 strain. As shown in Fig. 5, *MexD* gene expression was much higher in the mutant strains than in the wild-type strain, regardless of whether they were grown in high- or low-nutrient conditions. According to Purssell and Poole (10), the *MexCD-OprJ* efflux pump is quiescent in wild-type *P. aeruginosa* cells and does not contribute to intrinsic antibiotic resistance under standard laboratory conditions. However, mutations in the *nfxB* repressor lead to hyperexpression of the *MexCD-OprJ* efflux pump. Moreover, *MexD* gene expression was significantly higher (P < 0.05) under high-nutrient conditions containing (500 mg/l TOC; 48.89 ± 4.16-fold) than low-nutrient conditions (5 mg/l TOC; 8.01 ± 0.82-fold). Thus, less energy was needed for expression of the *MexCD-OprJ* efflux pump in *nfxB* mutants under low-nutrient conditions, which might have contributed to the reduction in fitness cost due to antibiotic resistance (24).

**Fig. 5.**
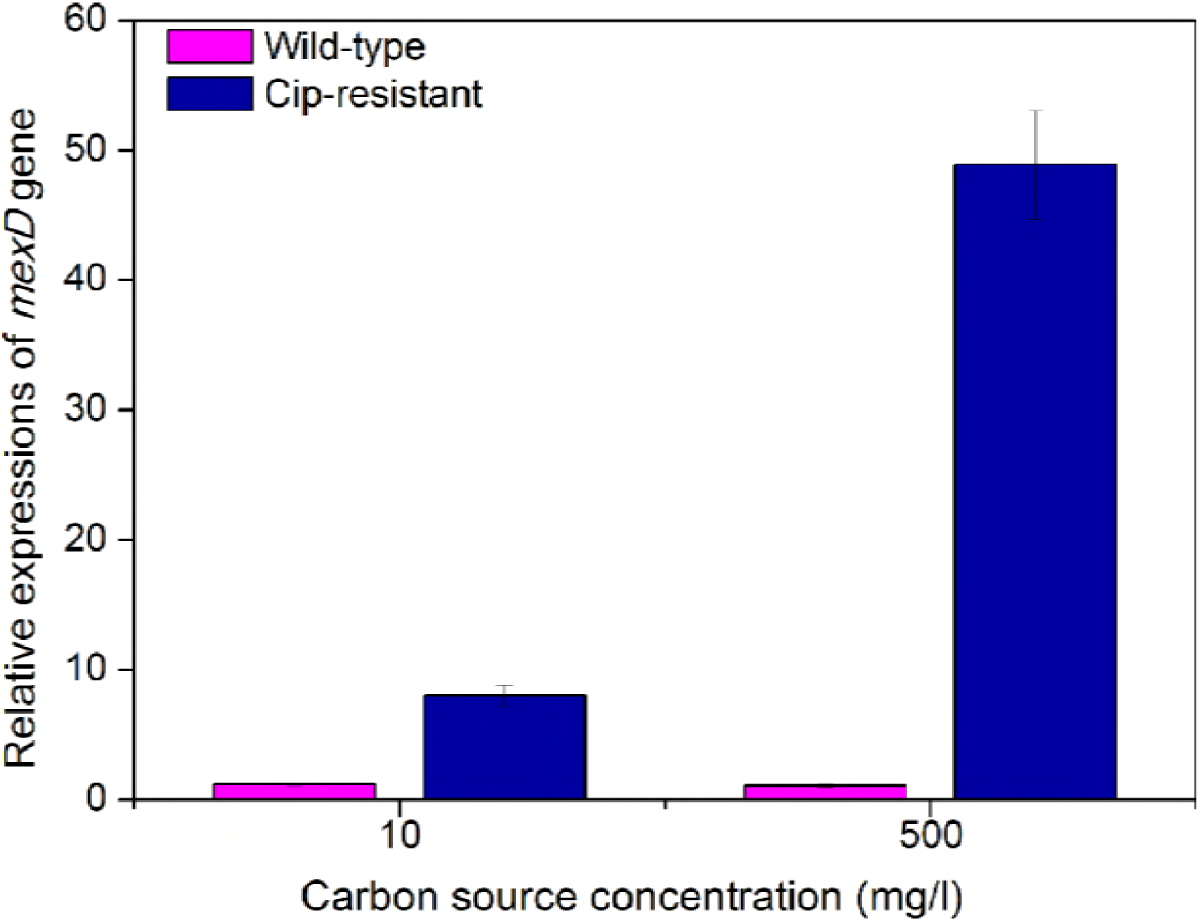
Relative amount of *MexD* gene expression under high (500 mg/l TOC) and low (10 mg/l) nutrient conditions by RT-qPCR. Levels of mRNA were normalized to that of the wild-type strain under low-nutrient conditions (set to 1.0).

### 3.3.2 *rpoS* gene regulation

Sigma factor (*rpoS*) is known to regulate the expression of hundreds of genes involved in adaption of bacteria in the stationary phase (25) and in osmotic conditions (26) and other stress environments (27). Therefore, in wild-type cells, the *rpoS* gene is induced in low-nutrient conditions, which inhibit their growth. As shown in Fig. 6, expression of the *rpoS* gene in the wild-type strain was significantly higher (P < 0.05) with 10 mg/l TOC (2.78 ± 0.29-fold) than with 500 mg/l TOC (1.02 ± 0.02-fold).

**Fig. 6.**
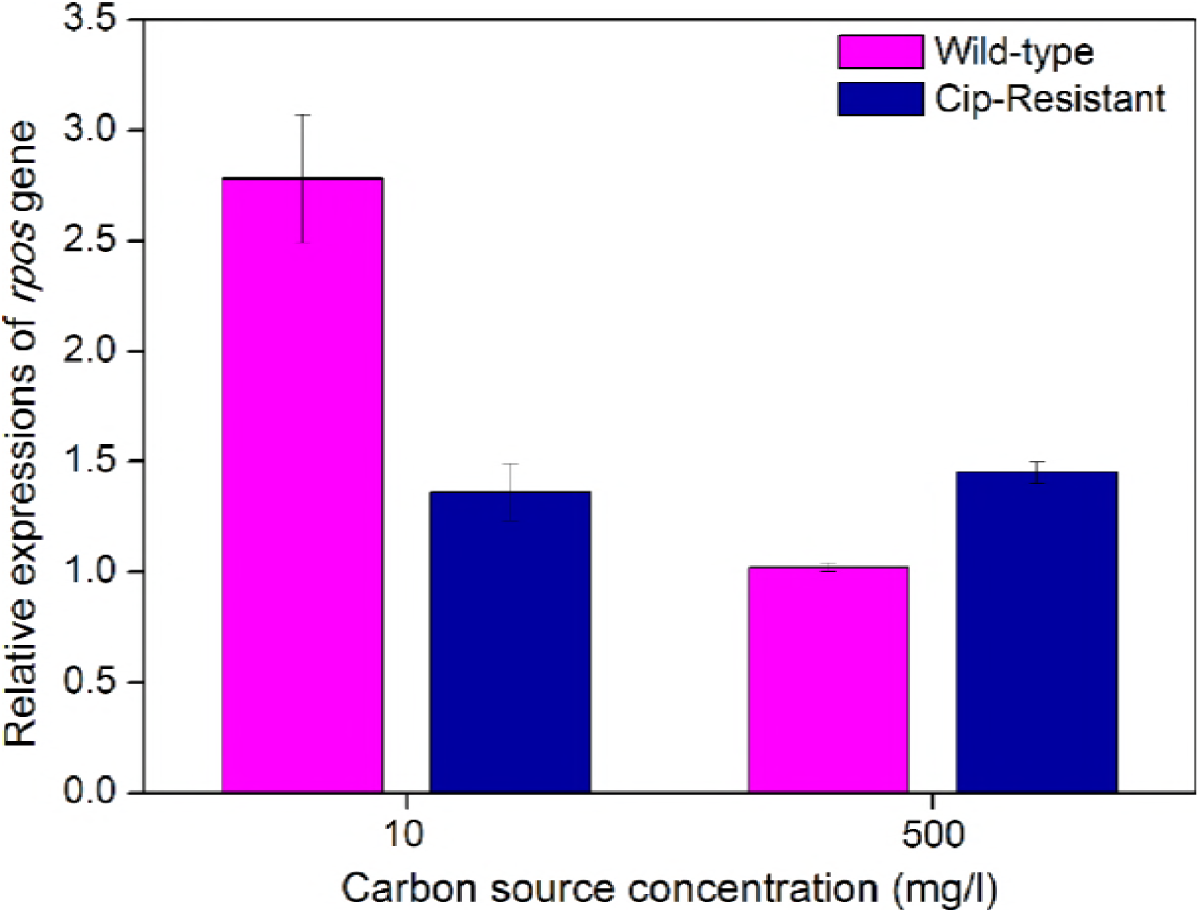
Relative amount of *rpoS* gene expression in *P. aeruginosa* PAO1 under high (500 mg/l TOC) and low (10 mg/l TOC) nutrient conditions by RT-qPCR. Levels of mRNA were normalized to that of the wild-type strain under nutrient-rich conditions (set to 1.0).

However, at a low nutrient level (10 mg/l TOC), *rpoS* gene expression in the resistant mutants (1.36 ± 0.13-fold) was significantly lower (P < 0.05) than that in the wild-type strain (2.78 ± 0.29-fold). Therefore, the inhibitory effect was less. The resistance mutants might show an advantage when competing with the wild-type strain. Similarly, in a previous study, Paulander et al. (28) found that the streptomycin resistance mutations *K42N* and *P90S* in ribosomal protein S12 impaired bacterial growth in a nutrient-rich medium, but that the mutants grew faster in poor nutrient conditions than the wild-type strain because the *rpoS* gene was not induced. Consequently, *rpoS* gene regulation might contribute to the reduced fitness cost of ciprofloxacin resistance under low-nutrient conditions.

### 3.3.3 Metabolic activity comparison

SOUR is an important indicator of microbial metabolism activity. In this study, the SOURs of ciprofloxacin-resistant strains were significantly lower (P < 0.05) than that of the wild-type strain at all nutrient levels (500, 5, and 0.5 mg/l TOC) (Fig. 7). For example, the SOURs of wild-type *P. aeruginosa* PAO1 were (7.19 ± 0.12) × 10^−12^, (3.16 ± 0.09) × 10^−12^, and (2.39 ± 0.07) × 10^−12^ mg O_2_/cells•h in the medium containing 500, 5, and 0.5 mg/l TOC, respectively. However, lower SOURs, i.e. (3.17 ± 0.05) × 10^−12^, (2.03 ± 0.05) × 10^−12^, and (1.80 ± 0.04) × 10^−12^ mg O_2_/cells•h, were observed for ciprofloxacin-resistant *P. aeruginosa* PAO1 at corresponding nutrient levels. These results indicated that the *nfxB* mutants had defects in metabolism. It is generally accepted that the acquisition of resistance is metabolically costly for bacteria (29). For example, Stickland et al. (11) found that a mutation in *nfxB* upregulated *MexCD*-*OprJ* expression, leading to global changes in *P. aeruginosa* PAO1 metabolism.

**Fig. 7.**
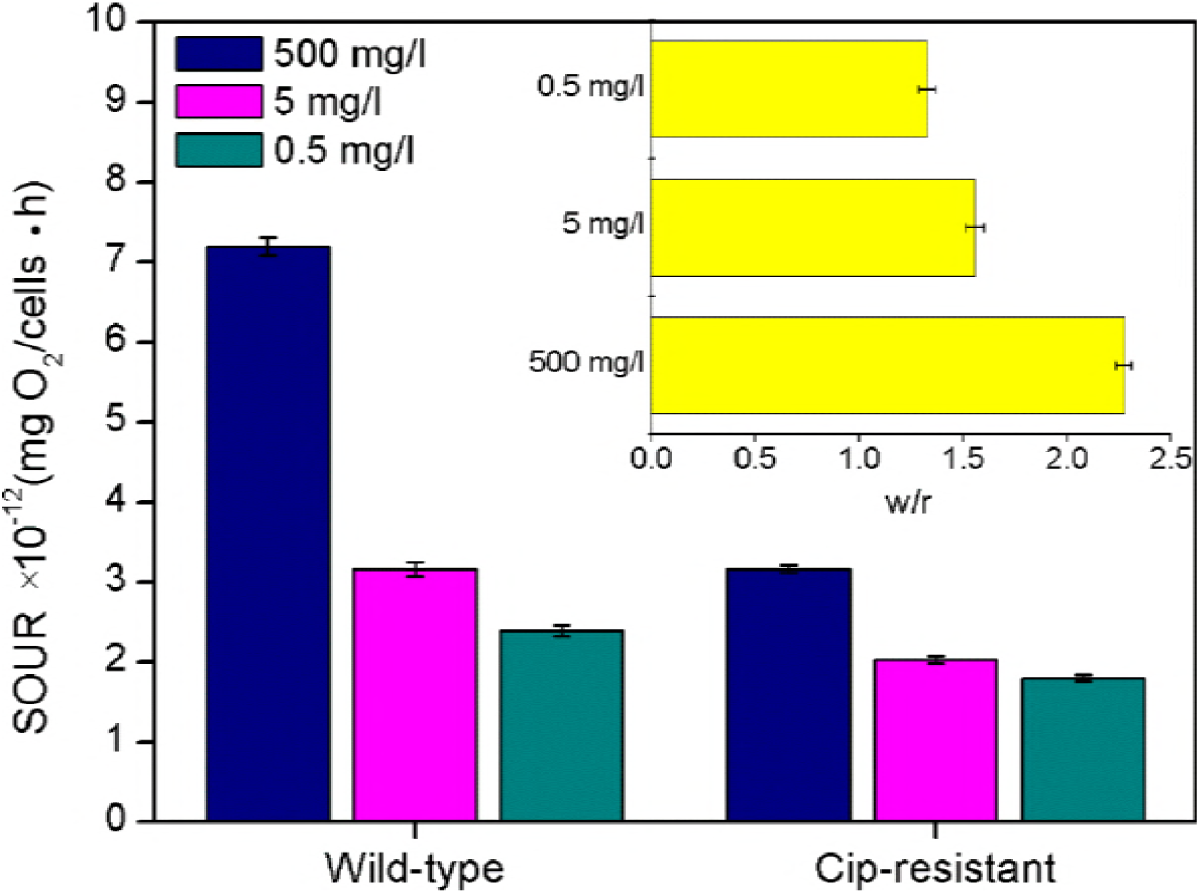
Respiratory rate of wild-type and ciprofloxacin resistant *P. aeruginosa* PAO1 at different nutrient levels (0.5, 5, and 500 mg/l TOC) by SOUR test. The data represent the average of three repeated independent experiments. The insets at the right top corner represent the ratios of SOURs between the wild-type and resistant strains.

Additionally, the difference in SOURs between the two competing strains was reduced with decreasing nutrient levels (Fig. 5). The differences in SOURs under low-nutrient conditions (5 mg/l TOC, 1.56 ± 0.04-fold; 0.5 mg/l TOC, 1.33 ± 0.04-fold) were significantly less (P < 0.05) than those at 500 mg/l TOC (2.27 ± 0.03-fold). This might be an explanation for the reduction in fitness cost at low nutrient levels (0.5 and 5 mg/l TOC).

## 4. Conclusion

Antibiotic resistance in environmental bacteria is becoming a public health problem. Efflux pumps play an important role in bacteria resistant to one or more antibiotics. This study shown that the ratio of the number of cells of two competing strains decreased and the average fitness of resistant mutants increased under low-nutrient conditions (0.05, 0.5, and 5 mg/l TOC), suggesting a reduction in fitness cost in the *nfxB* mutants in these cases. Some mechanisms, including those indicated by measures of *MexD* gene expression, SOURs, and *rpoS* gene regulation, were analyzed. *MexD* gene expression was shown to decrease in low-nutrient medium, meaning with lower energy consumption. In addition, *rpoS* gene expression levels were lower in the resistant mutants than in the wild-type strain in low-nutrient conditions, reducing the inhibitory effect of the gene product. Furthermore, the difference in SOURs between the two competing strains was reduced with decreasing nutrient levels. Therefore, low nutrient levels can reduce the fitness cost of ciprofloxacin resistance mediated by an efflux pump. In most natural environments such as source water, nutrient levels are low or even extremely low. Resistant bacteria can persist longer in these environments than in laboratory or clinical conditions; thus, antibiotic-resistant strains of bacteria in the environment are a reason for concern.

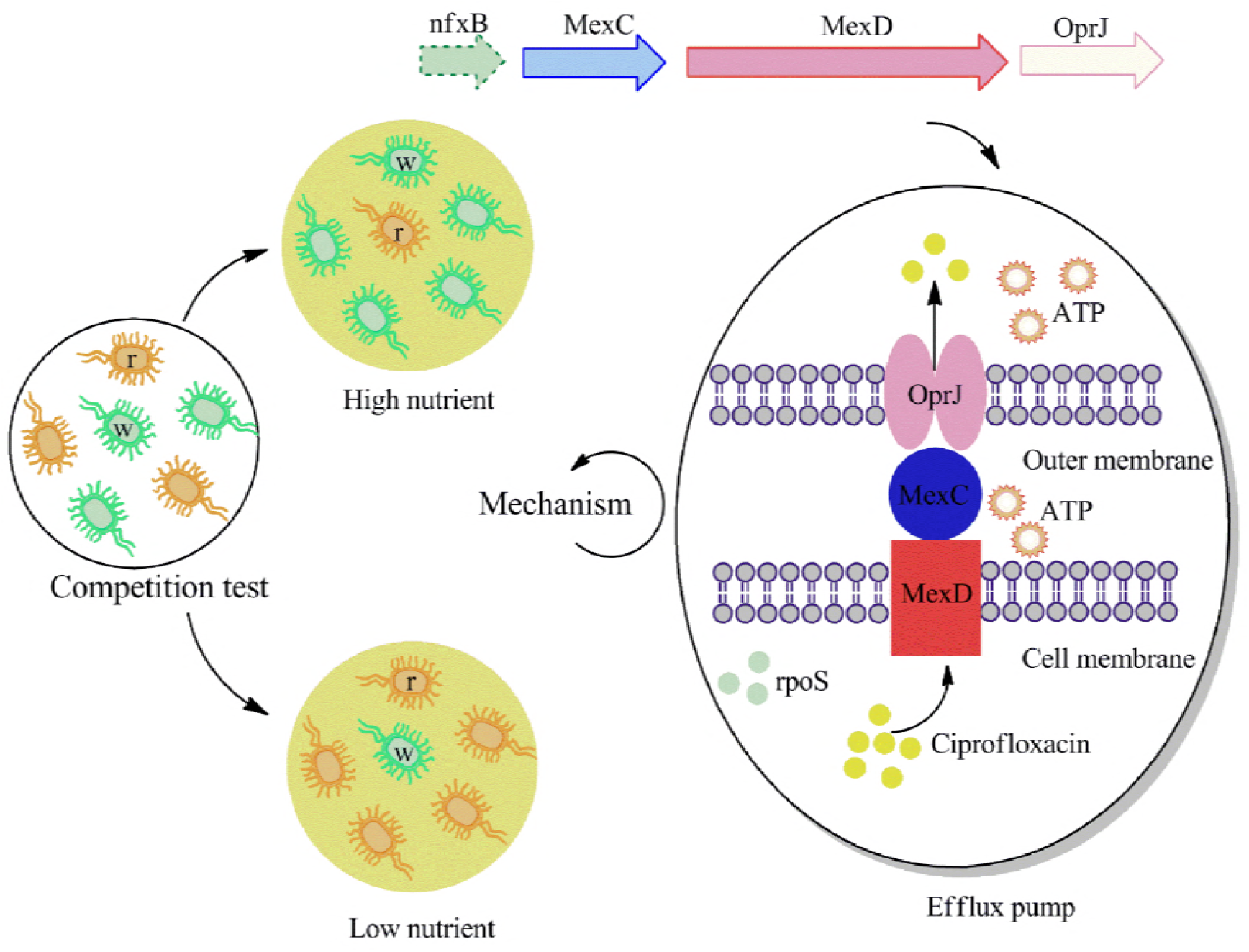

## Acknowledgements

This study was funded by the National Science Foundation of China (no. 51708534 and no. 51478450) and the National Key Research and Development Program of China-International collaborative project from the Ministry of Science and Technology (no. 2017YFE0107300).

## References

1. Baker-Austin C, Wright MS, Stepanauskas R, McArthur JV. 2006. Co-selection of antibiotic and metal resistance. Trends Microbiol. 14(4):176–182.

2. Delcour AH. 2009. Outer membrane permeability and antibiotic resistance. Biochimica et Biophysica Acta (BBA) - Proteins and Proteomics. 1794(5):808–816.

3. Wright GD. 2005. Bacterial resistance to antibiotics: Enzymatic degradation and modification. Adv Drug Delivery Rev. 57(10):1451–1470.

4. Lambert PA. 2005. Bacterial resistance to antibiotics: modified target sites. Adv Drug Delivery Rev. 57(10):1471–1485.

5. Webber MA, Piddock LJ. 2003. The importance of efflux pumps in bacterial antibiotic resistance. J Antimicrob Chemother. 51(1):9.

6. Poole K. 2001. Multidrug efflux pumps and antimicrobial resistance in *Pseudomonas aeruginosa* and related organisms. J Mol Microbiol Biotechnol. 3(2):255–264.

7. Hernando-Amado S, Blanco P, Alcalde-Rico M, Corona F, Reales-Calderón JA, Sánchez MB, Martínez JL. 2016. Multidrug efflux pumps as main players in intrinsic and acquired resistance to antimicrobials. Drug Resist Updates. 28(Supplement C):13–27.

8. Lister PD, Wolter DJ, Hanson ND. 2009. Antibacterial-resistant *Pseudomonas aeruginosa*: clinical impact and complex regulation of chromosomally encoded resistance mechanisms. Clin Microbiol Rev. 22(4):582–610.

9. Masuda N, Sakagawa E, Ohya S, Gotoh N, Tsujimoto H, Nishino T. 2000. Substrate specificities of *MexAB-OprM*, *MexCD-OprJ*, and *MexXY-oprM* efflux pumps in *Pseudomonas aeruginosa*. Antimicrob Agents Ch. 44(12):3322–3327.

10. Purssell A Poole K. 2013. Functional characterization of the *NfxB* repressor of the *MexCD-OprJ* multidrug efflux operon of *Pseudomonas aeruginosa*. Microbiology. 159(10):2058–2073.

11. Stickland HG, Davenport PW, Lilley KS, Griffin JL, Welch M. 2010. Mutation of *nfxB* Causes Global Changes in the Physiology and Metabolism of *Pseudomonas aeruginosa*. J Proteome Res. 9(6):2957–2967.

12. Andersson DI, Hughes D. 2010. Antibiotic resistance and its cost: is it possible to reverse resistance? Nat Rev Microbiol. 8(4):260–271.

13. Hirai K, Suzue S, Irikura T, Iyobe S, Mitsuhashi S. 1987. Mutations producing resistance to Norfloxacion in *Pseudomonas aeruginosa*. Antimicrob Agents Ch. 31(4):582–586.

14. Jeannot K, Elsen S, Köhler T, Attree I, Van Delden C, Plésiat P. 2008. Resistance and virulence of *Pseudomonas aeruginosa* clinical strains overproducing the *MexCD-OprJ* efflux pump. Antimicrob Agents Ch. 52(7):2455–2462.

15. Olivares J, Alvarez-Ortega C, Linares JF, Rojo F, Köhler T, Martínez JL. 2012. Overproduction of the multidrug efflux pump *MexEF-OprN* does not impair *Pseudomonas aeruginosa* fitness in competition tests, but produces specific changes in bacterial regulatory networks. Environ Microbiol. 14(8):1968–1981.

16. Björkman J, Nagaev I, Berg O, Hughes D, Andersson DI. 2000. Effects of environment on compensatory mutations to ameliorate costs of antibiotic resistance. Science. 287(5457):1479–1482.

17. Gifford DR, Moss E, MacLean RC. 2016. Environmental variation alters the fitness effects of rifampicin resistance mutations in *Pseudomonas aeruginosa*. Evolution. 70(3):725–730.

18. Lv L, Jiang T, Zhang SH, Yu X. 2014. Exposure to mutagenic disinfection byproducts leads to increase of antibiotic resistance in *Pseudomonas aeruginosa*. Environ Sci Technol. 48(14):8188–8195.

19. Sander P, Springer B, Prammananan T, Sturmfels A, Kappler M, Pletschette M, Böttger EC. 2002. Fitness cost of chromosomal drug resistance-conferring mutations. Antimicrob Agents Ch. 46(5):1204–1211.

20. Song T, Park Y, Shamputa IC, Seo S, Lee SY, Jeon HS, Choi H, Lee M, Glynne RJ, Barnes SW. 2014. Fitness costs of rifampicin resistance in *Mycobacterium tuberculosis* are amplified under conditions of nutrient starvation and compensated by mutation in the β′ subunit of RNA polymerase. Mol Microbiol. 91(6):1106–1119.

21. Zhang SH, Yu X, Guo F, Wu ZY. 2011. Effect of interspecies quorum sensing on the formation of aerobic granular sludge. Water Sci Technol. 64(6):1284–1290.

22. Livak KJ Schmittgen TD. 2001. Analysis of Relative Gene Expression Data Using Real-Time Quantitative PCR and the 2^−ΔΔCT^ Method. Methods. 25(4):402–408.

23. Andersson DI. 2006. The biological cost of mutational antibiotic resistance: any practical conclusions? Curr Opin Microbiol. 9(5):461–465.

24. Schweizer HP. 2003. Efflux as a mechanism of resistance to antimicrobials in *Pseudomonas aeruginosa* and related bacteria: unanswered questions. Genet Mol Res. 2(1):48–62.

25. Hengge-Aronis R. 1993. Survival of hunger and stress: The role of *rpoS* in early stationary phase gene regulation in *E. coli.* Cell. 72(2):165–8.

26. Shiroda M, Pratt ZL, Wong AC, Kaspar CW. 2014. *RpoS* impacts the lag phase of *Salmonella enterica* during osmotic stress. FEMS Microbiol Lett. 357(2):195.

27. O’Neal CR, Gabriel WM, Turk AK, Libby SJ, Fang FC, Spector MP. 1994. *RpoS* is necessary for both the positive and negative regulation of starvation survival genes during phosphate, carbon, and nitrogen starvation in *Salmonella typhimurium*. J Bacteriol. 176(15):4610.

28. Paulander W, Maisnier-Patin S, Andersson DI. 2009. The fitness cost of streptomycin resistance depends on rpsL mutation, carbon source and *RpoS* (σ^S^). Genetics. 183(2):539–546.

29. Bhargava P, Collins J. 2015. Boosting bacterial metabolism to combat antibiotic resistance. Cell Metab. 21(2):154–155.

